# Modeling Atrial Fibrillation through Intermittent Tachypacing-Induced Remodeling in hiPSC-Derived Atrial Cardiomyocytes and Atrial Fibroblast

**DOI:** 10.1101/2025.05.23.655123

**Authors:** Paul Kozlowski, Kalai Mangai Muthukumarasamy, Amrish Deshmukh, Andre Monteiro Da Rocha, Hakan Oral

**Author notes:** Equal contribution (Paul Kozlowski, Kalai Mangai Muthukumarasamy). Contact Author: Hakan Oral, MD., Cardiovascular Center, SPC 5853, 1500 East Medical Center Dr, Ann Arbor, MI 48109-5853.

## Abstract

**Background:** Human *in vitro* models for atrial fibrillation (AF) are limited. Human-induced pluripotent stem cell-derived atrial cardiomyocytes (hiPSC-aCMs) provide a valuable tool to study AF pathophysiology by facilitating *in vitro* modeling.

**Objectives:** To investigate the effects of an intermittent tachypacing protocol (ITPP) in matured hiPSC-aCMs co-cultured with human atrial cardiac fibroblast (haCF) to mimic AF-associated electroanatomical phenotypes.

**Methods and Results:** hiPSC-aCMs were cultured alone or co-cultured with haCFs at 90/10 and 70/30 ratios. ITPP was applied through field stimulation, and optical mapping assessed action potentials (APs) and calcium transients (CaTs). Immunostaining was performed to quantify pro-fibrotic biomarkers (Collagen III and TGFβ1). ITPP led to increased spontaneous AP frequency (Δ=+31±7%, P<0.0001) and reduced AP duration at 80% repolarization (APD_80%_; Δ=-15±4%, P=0.001). Additionally, the upstroke slope (Δ=-41±11%, P=0.001) and amplitude (dF/F_0_; Δ=-51±13%, P<0.001) of intracellular CaT were significantly reduced. Co-culture at the 70/30 hiPSC-aCM/haCF ratio, showed a >100-fold increase in Collagen III expression (P<0.0001), diminished excitability (ΔHz=-61±6%, P<0.0001), prolonged ΔAPD_80%_ (Δ=+130±10%, P<0.0001), prolonged AP triangulation (ΔAPD_Tri_=+143±13%, P<0.0001), reduced upstroke slope (Δ=-66±6%, P<0.0001), conduction block (Δ=-52±18%, P=0.0260), and diminished intracellular calcium handling (upstroke slope Δ=-50±8%, P<0.0001; ΔdF/F0=-34±9%, P=0.0003). Finally, the application of ITPP to the 70/30 co-culture model recapitulated an AF-mediated phenotype (ΔHz=+25±8%, P=0.02; ΔAPD_80%_=-16±6%, P=0.01) while introducing conduction block (ΔCV_100/0 vs 70/30_= −27±15%; P=0.0005).

**Conclusions:** Co-cultures of matured hiPSC-aCMs and haCFs exhibited structural and electrophysiological remodeling, including conduction abnormalities, mirroring key AF mechanisms. This model holds potential for patient-specific therapies and drug discovery.

Graphical Abstract
Human iPSC-Model for Atrial Fibrillation

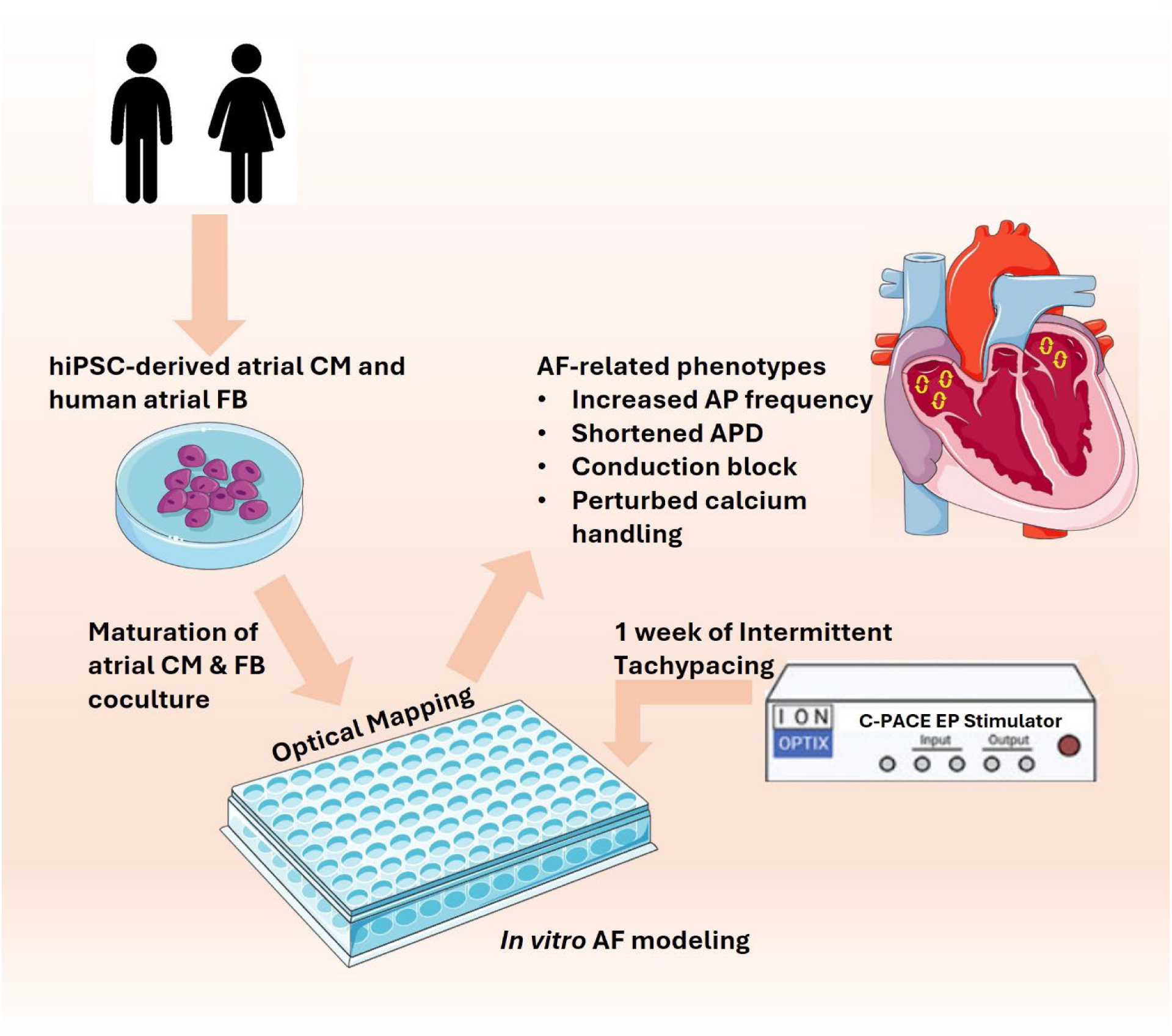

## INTRODUCTION

A variety of animal models have been utilized to study AF, ranging from small animal models, such as mice, rats, guinea pigs, and rabbits, to larger animal models including goats, canines, sheep, pigs, and horses (1). Computational models of AF integrate data from clinical studies, *in vitro* experiments, and animal studies to provide insights into risk prediction, underlying mechanisms, and potential therapeutic strategies (2,3). Challenges in extrapolating findings from these models to humans, compounded by limitations in current model organisms and experimental methods that do not fully capture the complexities of human AF mechanisms, further complicate the translation of scientific discoveries into viable clinical treatments (4). In this context, human pluripotent stem cells (hPSCs) have emerged as a valuable tool for personalized medicine, drug discovery (5–10), and the study of cardiac genetics and diseases (11–15).

In the present study, we utilized a well-studied perinatal stem cell derived extracellular matrix (ECM) (16) that promotes the development of highly organized sarcomeres, elevated levels of cardiac troponin I (cTnI), enhanced mitochondrial distribution, and improved electrophysiological function, thereby approximating the characteristics of mature CMs (17). Furthermore, to address the inherent heterogeneity within the cardiac cell population (18,19), we implemented a retinoic acid (RA)-based atrial-specific monolayer differentiation protocol (20). This was followed by the purification of the cardiomyocyte population, which was subsequently matured to facilitate downstream intermittent pacing experiments to model AF in mature hiPSC-aCM-haCF cocultures.

## METHODS

### Intermittent Tachypacing of hiPSC-aCM and hiPSC-aCM/haCF Co-Culture

Human iPSC-aCMs differentiated (21) from DF19-9-11T human iPSC line (WiCell Research Institute) and matured as previously described (17,22) were co-cultured with varying percentages of haCFs (Lonza, #CC-2903): 100%, 90%, or 70%. For pacing experiments, the cells were plated in 8-well plates coated with MatrixPlus (StemBioSys) and fitted with a polydimethylsiloxane silicone sheeting (Specialty Manufacturing, Inc, Saginaw, MI) imprinted with a circle (3.14 cm^2^) of free space to restrict the seeding area of the cells. The cells were subjected to chronic electrical field pacing using a C-PACE EP Cell Culture Stimulator Bank (IONOptix) coupled with C-Dish™ 8-well carbon electrodes (IONOptix). An intermittent tachypacing protocol was established by applying electrical field stimulation at 20 V/cm with 0.5 ms pulse duration at frequency cycles at 1.5 Hz for 45 minutes followed by 5 Hz for 15 minutes, repeated hourly for a duration of 7 days. This study was conducted in accordance with the Human Pluripotent Stem Cell Research Oversight (HPSCRO #0147) ethical approval. The research presented in this manuscript is the subject of a pending patent application **63/761,636** (USPTO), filed by Drs. Oral and Monteiro da Rocha.

### High-Throughput Optical Mapping

Optical mapping was performed using a CARTOX^TM^ device (StemBioSys) at 37 °C. For mapping of calcium transients (CaTs) hiPSC-aCM monolayers were pre-loaded with the GCaMP6f, as previously described (17). Mapping of CaTs in hiPSC-aCM/haCF co-culture preparations was facilitated using the Calbryte™ 520 AM intracellular calcium indicator (20651; AAT Bioquest). Action potentials were recorded using the FluoVolt probe (F10488; Life Technologies). Following a 30-minute incubation, monolayers were washed with HBSS with calcium and magnesium, then incubated at 37 °C for 1 hour prior to optical mapping. Data analysis was conducted using StemBioSys software.

Detailed description of the methods can be found in the **Supplementary material**.

## RESULTS

### Tachycardia-mediated Phenotype of APs in ITPP-treated hiPSC-aCM syncytia

Optical mapping of APs (Figure 1A) revealed that ITPP-treated hiPSC-aCM syncytia exhibited a significantly higher frequency of spontaneous APs relative to control (P<0.0001; Figure 1B). Fridericia-corrected APD_80%_ (P=0.001; Figure 1C) and AP triangulation (P=0.01; Figure 1D) were both shortened in ITPP-treated cells, suggesting a reduction in phase 3 repolarization. The upstroke slope of the AP, a measurement of phases 0/1, was not affected by ITPP (P=0.78; Figure 1E). Interestingly, the conduction velocity of the AP was significantly faster following ITPP (P<0.001; Figure 1F).

**Figure 1.**
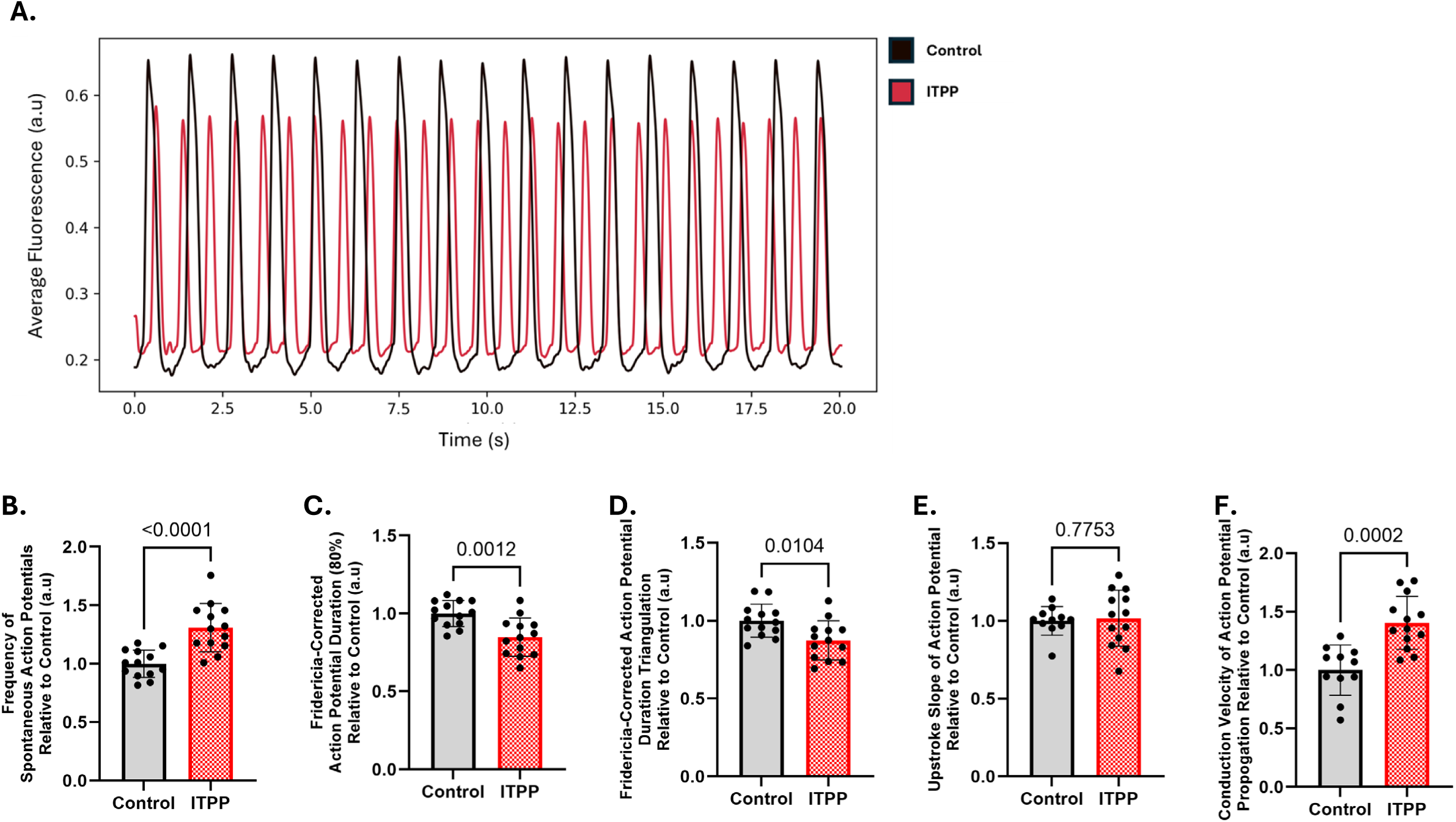
Functional electrical remodeling of hiPSC-aCM syncytia submitted to ITPP. (A) Time-sequence plot of representative action potential traces of hiPSC-aCM syncytia of control (black) and ITPP (red). (B) Paced hiPSC-aCM syncytia exhibited an increased frequency of spontaneous APs relative to control (Δ=+31±7%, p<0.0001) alongside reductions in Fridericia-corrected AP duration (80%) (Δ=-15±4%, P=0.0012) (C) and AP triangulation (D) (Δ=-13±5%, P=0.0104), with preserved upstroke slope (E) and significantly faster conduction velocity (F) (Δ=+41±9%, P=0.0002).

### ITPP alters calcium handling similar to observed in AF

To assess the effects of ITPP on intracellular calcium handling, hiPSC-aCM syncytia were further evaluated by optical mapping to study CaTs (Figure 2A). Foremost, the variability of spontaneous calcium release events was measured using the standard deviation of frequency of spontaneous CaTs. While the baseline frequency of spontaneous CaTs was not significantly different between ITPP-treated hiPSC-aCM syncytia relative to controls (P=0.16; Figure 2B), there was a significant increase (Δ = +332±90%) in the variability of calcium release events following ITPP (P<0.001; Figure 2C).

**Figure 2.**
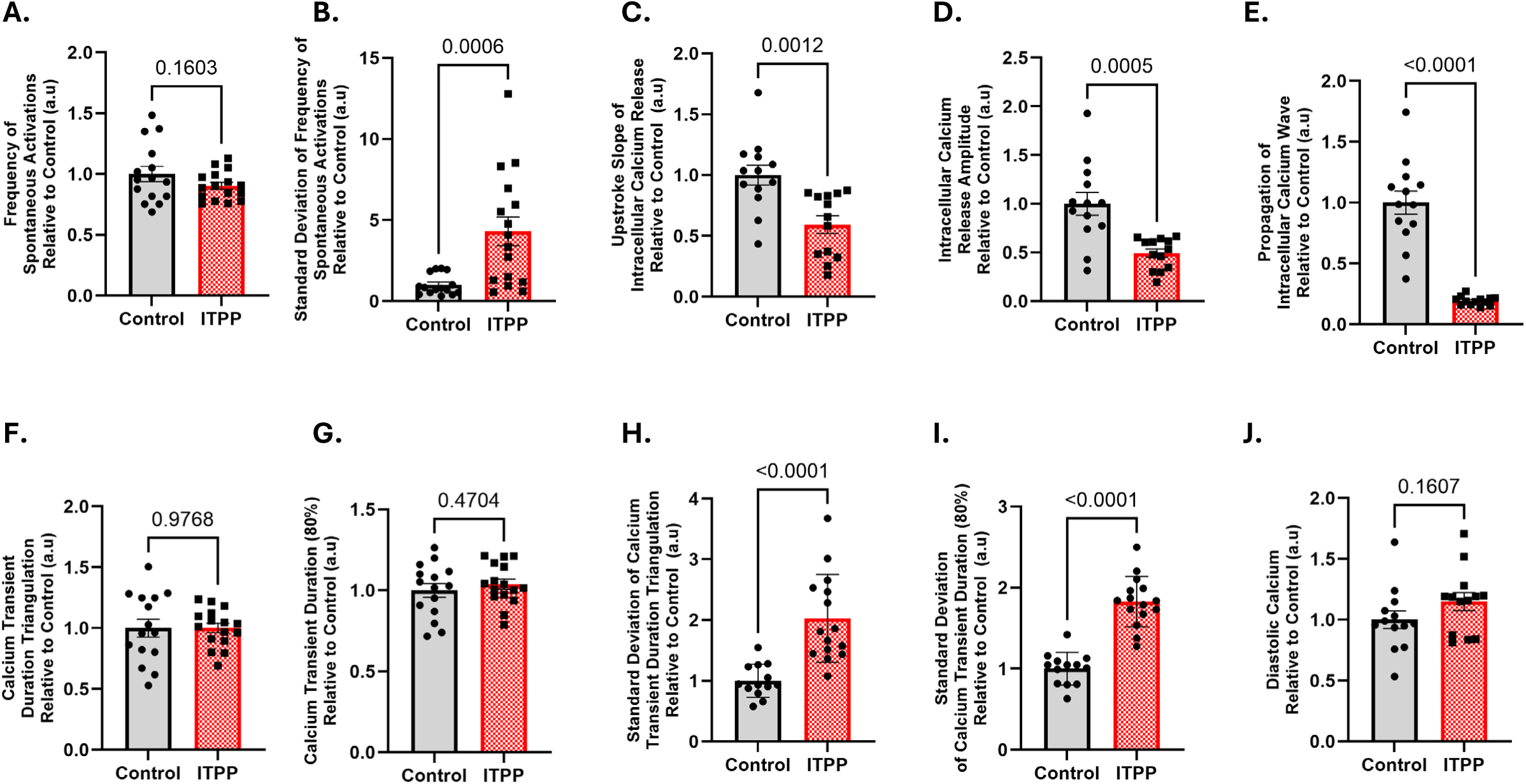
Intracellular calcium handling alterations of hiPSC-aCM syncytia subjected to ITPP. (A) ITPP did not influence the frequency of spontaneous calcium release but markedly increased the variability of calcium release events (Δ=+332±90%, P=0.0006) (B). (C) ITPP decreased both the upstroke slope (Δ=-41±11%, P=0.0012) and (D) amplitude of intracellular calcium release (Δ=-51±13%, P=0.0005) with slowed propagation speed of the calcium wave (E) (Δ=-81±10%, P<0.0001). No change in average speed of SR calcium reuptake (F) and in CaTD_80%_ duration (G) was affected by ITPP. (H-I) The beat-to-beat variability of both CaTD_Tri_ (Δ=+103±20%, P<0.0001) and CaTD_80%_ (Δ=+83±10%, P<0.0001) was significantly increased. (J) The diastolic calcium level following the CaT remained unchanged following ITPP.

The upstroke slope of the CaT measures the speed of calcium release from the sarcoplasmic reticulum. Compared to controls, ITPP-treated hiPSC-aCM syncytia exhibited a significantly slower upstroke slope of calcium release (P=0.001; Figure 2D), suggesting impaired SR function. Calcium release amplitude was also significantly diminished in ITPP-treated cells (P<0.001; Figure 2E), and the propagation speed of the intracellular calcium wave across the monolayer was significantly slower (P<0.001; Figure 2F).

The duration of intracellular calcium reuptake into the sarcoplasmic reticulum was assessed using a CaT triangulation measure; nevertheless, we did not observe any significant, relative change to the average value of the CaTD_Tri_ (P=0.98; Figure 2G). There was also no change observed in the average duration at 80% decay of the CaT between the groups (P=0.47; Figure 2H). However, there was a significant increase in the beat-to-beat variability in SR reuptake duration (P<0.0001; Figure 2I) in ITPP-treated hiPSC-aCMs compared to controls, alongside an increase in the standard deviation of CaTD_80%_ (P<0.0001; Figure 2J) consistent with our observation of increased variability in the frequency of spontaneous CaT events. There was no significant change in diastolic calcium levels (F0) between groups immediately following our 7-day ITPP protocol (P=0.16; Figure 2K).

Optical mapping was also performed before and after administration of Arrythmia Inducibility Test (AIT; Figure 3A) in control and ITPP-treated hiPSC-aCM syncytia. Comparison of the difference in average baseline fluorescence during and after AIT serves to assess the change in diastolic calcium levels following burst tachypacing. In post-rest potentiation (PRP) of CaTs, ITPP-treated hiPSC-aCMs exhibited a reduced upstroke slope (P<0.001; Figure 3B) and decreased amplitude (P<0.001; Figure 3C) relative to controls. ITPP-treated hiPSC-aCM syncytia also exhibited a significantly diminished reconstitution of diastolic calcium post-AIT, (P<0.001; Figure 3D) consistent with observed alterations to PRP CaT dynamics.

**Figure 3.**
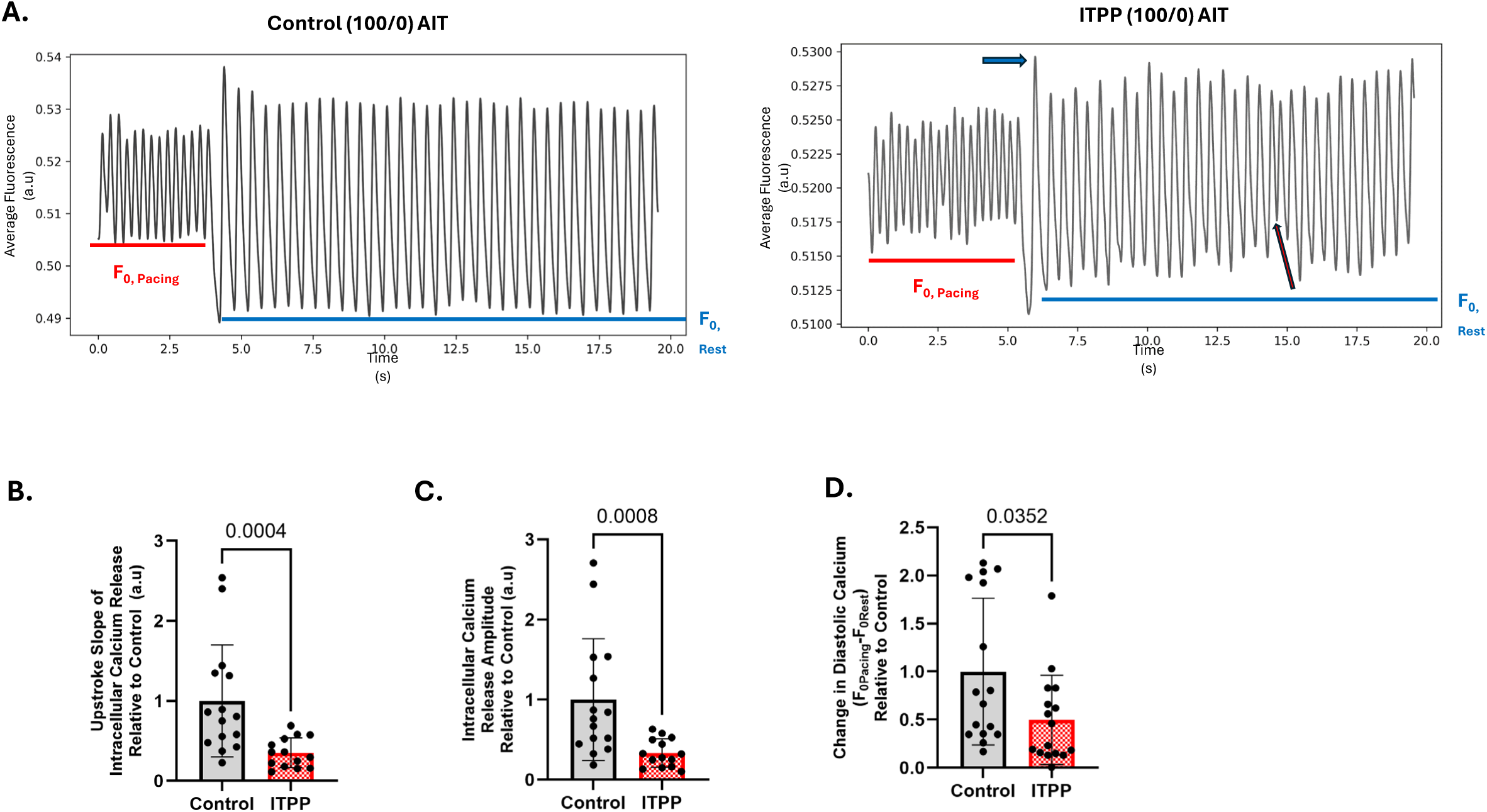
Arrhythmia inducibility test reveals calcium overload susceptibility in hiPSC-aCM syncytia following ITPP. (A) Representative traces of AIT stimulation in Control and ITPP hiPSC-aCM syncytia demonstrate transition between AIT pacing and rest. The post-rest potentiation calcium transient (PRP-CaT) marking the transition from pacing to rest is labeled with a blue arrow; a premature calcium release event is labeled with a red arrow. Diastolic calcium levels during AIT pacing (red line) and rest (blue line) are indicated. (B) The PRP-CaT, marking calcium flux during transition from AIT pacing to rest, presented decreased upstroke slope of intracellular calcium release (Δ=-65±19%; P=0.0004), (C) decreased intracellular calcium release amplitude (Δ=-67±20%; P=0.0008), and (D) reduced recovery of baseline calcium levels after AIT cessation (Δ=-50±22%, P=0.0352).

### hiPSC-aCM-haCF co-culture recapitulates characteristics of AF-associated atrial fibrosis and conduction slowing/block

Co-culture of purified and matured hiPSC-aCM with haCFs was conducted at different seeding density ratios of 100/0, 90/10, and 70/30 to explore fibroblast-induced electrophysiological and calcium-handling changes. Cocultures maintained the seeding ratios following 7 days of culture (100/0 cTnT:TE7: 0.96±0.03:0.01±0.001; 90/10 cTnT:TE7: 0.84±0.03:0.15±0.01; 70/30 cTnT:TE7: 0.72±0.03: 0.34±0.03; Figure 4A-B). To verify pro-fibrotic effects of haCF culture, Collagen III expression was measured using immunocytochemistry. The 70/30 ratio (hiPSC-aCM/haCF) exhibited the highest expression of collagen III, which was fibroblast-dependent (P<0.0001; Figure 4C-D). Expression of the upstream cytokine TGFβ1 was also significantly increased in ratios consistent with increased collagen III expression (P=0.0026; Figure 4E-F).

**Figure 4.**
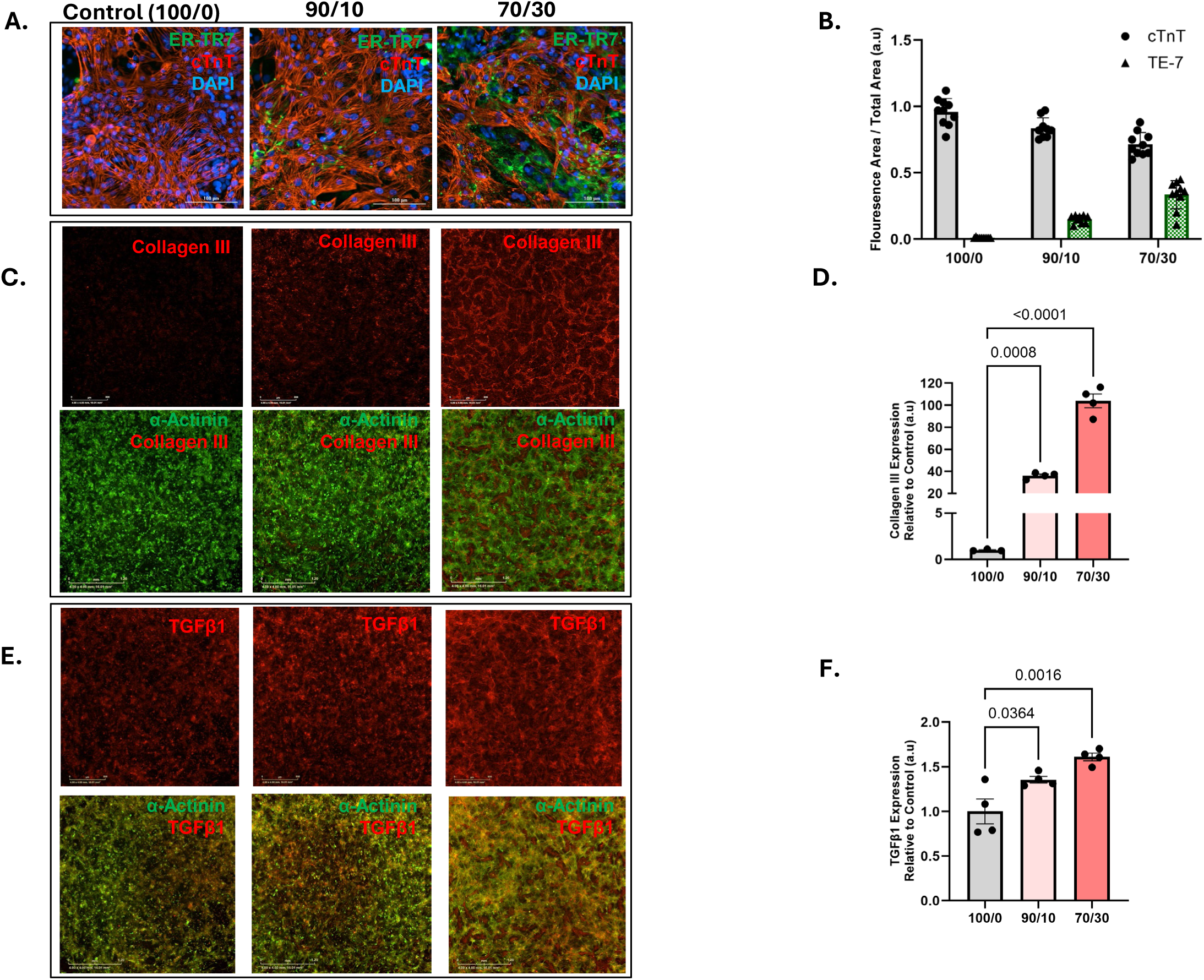
hiPSC-aCMs and haCFs co-culture induces atrial fibrosis. (A) Representative micrographs of cTnT/TE7 immunostaining in control (100/0) and co-culture (90/10, 70/30), and its (B) Fluorescence quantification after 7 days of co-culture with cTnT:TE7 for 100/0 (0.96±0.03 to 0.01±0.001), 90/10 (0.84±0.03 to 0.15±0.01) and 70/30 (0.72±0.03 to 0.34±0.03). (C) Representative micrograph panels of immunostaining for Collagen III (C) and TGFβ1 (E) with α-actinin labeling to highlight the cardiomyocyte fraction in the preparation. (D) Quantification shows a proportional increase with haCFs ratio in Collagen III (D) and TGFβ1 (F).

A previous report (23) demonstrated that the effect of fibroblast co-culture on the electrophysiology of hiPSC-aCMs is ratio-dependent. Therefore, we subjected hiPSC-aCM-haCF co-culture preparations to voltage and calcium optical mapping. Voltage mapping (Figure 5A) revealed fibroblast-dependent decrease to spontaneous AP frequency (P<0.0001; Figure 5B) with concomitant prolongation of both the Fridericia-corrected AP duration at 80% repolarization (P<0.0001; Figure 5C) and Fridericia-corrected APD triangulation (P<0.0001; Figure 5D) at 70/30 ratio only. Substantially, the upstroke slope of the AP (P<0.0001; Figure 5E) and conduction velocity of the AP (P<0.0001; Figure 5F) were also decreased at 70/30 ratio.

**Figure 5.**
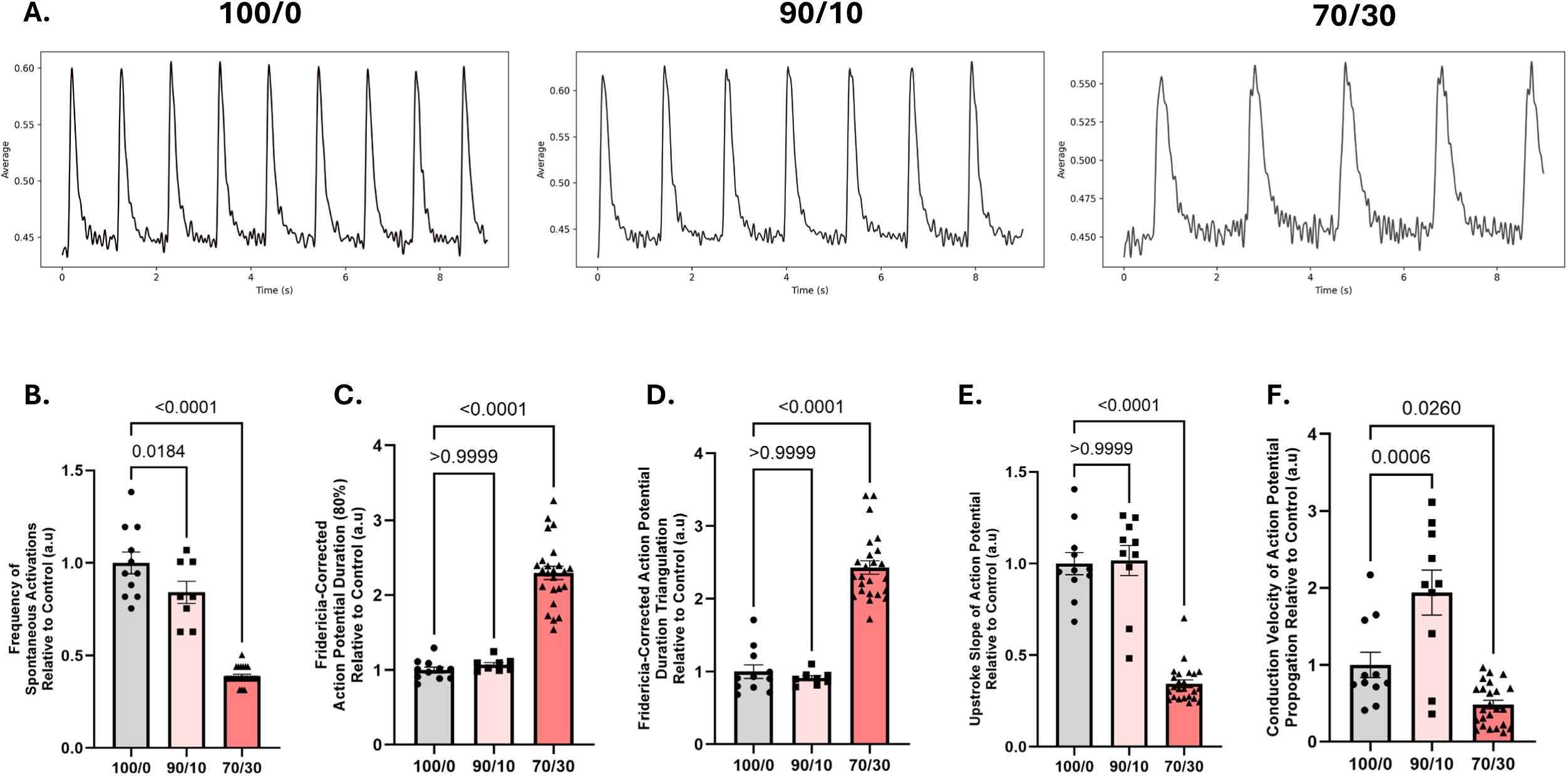
Fibrosis alters electrophysiology of hiPSC-aCM co-culture. (A) Representative time-sequence plots of action potential traces of control (100/0) and hiPSC-aCM/haCF co-culture at 90/10 and 70/30 ratios. (B) Spontaneous activation frequency decreased with increasing fibroblast proportion (90/10: Δ=-16±9%, P=0.0184; 70/30: Δ=-61±6%, p<0.0001). Co-culture with 30% haCFs further achieved prolongation of the Fridericia-corrected AP duration (80%) (C) (70/30: Δ=+130%±10%, P<0.0001), prolongation of the Fridericia-corrected AP triangulation (D) (70/30: Δ=+143±13%, P<0.0001), and decrease in the upstroke slope of the AP (E) (70/30: Δ=-66±6%, P<0.0001) with conduction slowing (F) (70/30: Δ=-52±18%, P=0.0260).

Evaluation of co-culture intracellular calcium transients also demonstrated ratio-dependent changes observed to functional electrophysiology, particularly at 70/30 conditions (Figure 6A). Co-culture did not affect the average frequency of intracellular calcium release events (P=0.65; Figure 6B), however it did prolong the both duration of 80% decay of the CaT (P<0.001; Figure 6C), and the duration of SR calcium reuptake (P=0.002; Figure 6D) in a ratio-dependent manner. Interestingly, 90/10 coculture presented a modest, significant decrease to diastolic calcium, however this was not recapitulated in 70/30 preparations (P=0.77; Figure 6E).

**Figure 6.**
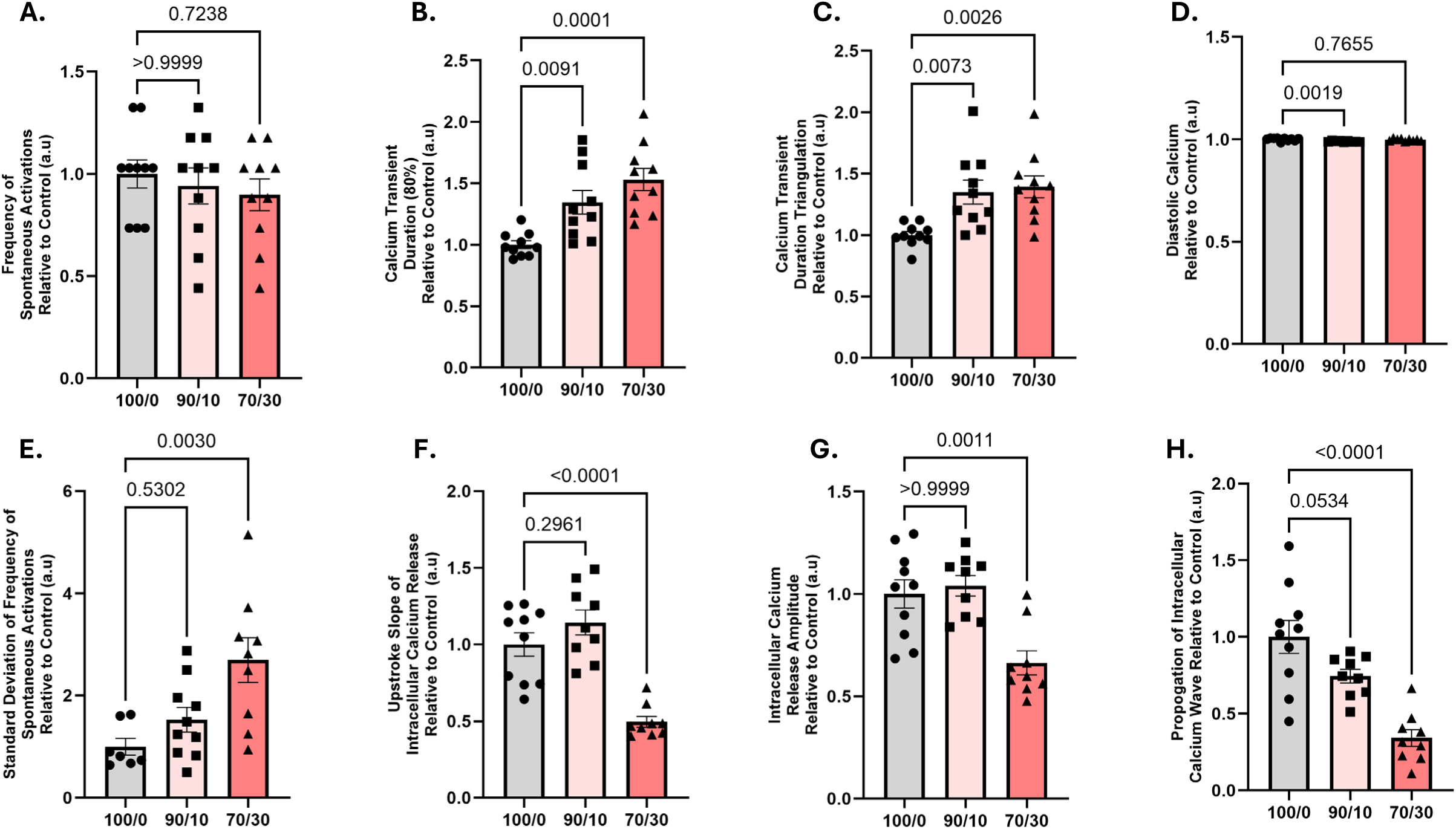
Fibrosis disrupts intracellular calcium handling of hiPSC-aCM co-culture. (A) No significant difference in the frequency of spontaneous calcium release events. (B) Prolongation of both the calcium transient duration (80%) (Δ=+53±10%, P=0.0002), and (C) duration of SR calcium reuptake was observed in ratio-dependent manner, most prominently at 70/30 co-culture (Δ=+39±9%, P=0.0022). (D) The diastolic calcium was decreased in 90/10 preparations (Δ=-1±0.25%, P=0.0028) without an effect at 70/30. (E) Significant increase in calcium release event variability at 70/30 co-culture (Δ=+170±47%, P=0.0030) with reduced the upstroke slope (F) (Δ=-50±8%, P<0.0001), and amplitude (G) (Δ=-34±9%, P=0.0003) of intracellular calcium release. (H) Decrease in intracellular propagation of the calcium wave by 70/30 co-culture (Δ=-66±12%, P<0.0001).

Calcium release events became markedly more variable at 70/30 ratio (P=0.0030; Figure 6F), with diminished intracellular calcium upstroke slope (P<0.0001; Figure 6G) and intracellular calcium release amplitude (P=0.0003; Figure 6H). The propagation of the intracellular wave was significantly slower in 70/30 preparations (P<0.0001; Figure 6I).

Given the similarities in the functional electrophysiology and calcium handling of 70/30 coculture to atrial fibrosis – a critical structural component of AF propagation, we hypothesized that coculture preparations subjected to electrical remodeling by ITPP would present conduction block analogous to this known proarrhythmic substrate. Voltage mapping of ITPP-treated co-cultures at 100/0 (Control; F-ITPP 100/0) and 70/30 (ITPP; F-ITPP 70/30) ratios verified that 70/30 preparations recapitulated tachycardia-mediated electrical remodeling observed in 100/0 iPSC-aCM syncytia (Figure 7A-C), including increased frequency of spontaneous activations (P=0.002; Figure 7D) and shortening of the Fridericia-corrected APD_80%_ (P=0.014; Figure 7E). The magnitude of change in frequency (P=0.42 Figure 7D) or APD80% (P=0.86; Figure 7E) was preserved between F-ITPP 70/30 or F-ITPP 100/0 conditions. Critically, F-ITPP 70/30 preparations did not demonstrate facilitated increase in the conduction velocity of the AP (P<0.001; Figure 7F). Rather, conduction velocity was significantly decreased by inclusion of fibroblasts in F-ITPP 70/30 preparations in relation to F-ITPP 100/0 (P=0.0001; Figure 7F).

**Figure 7.**
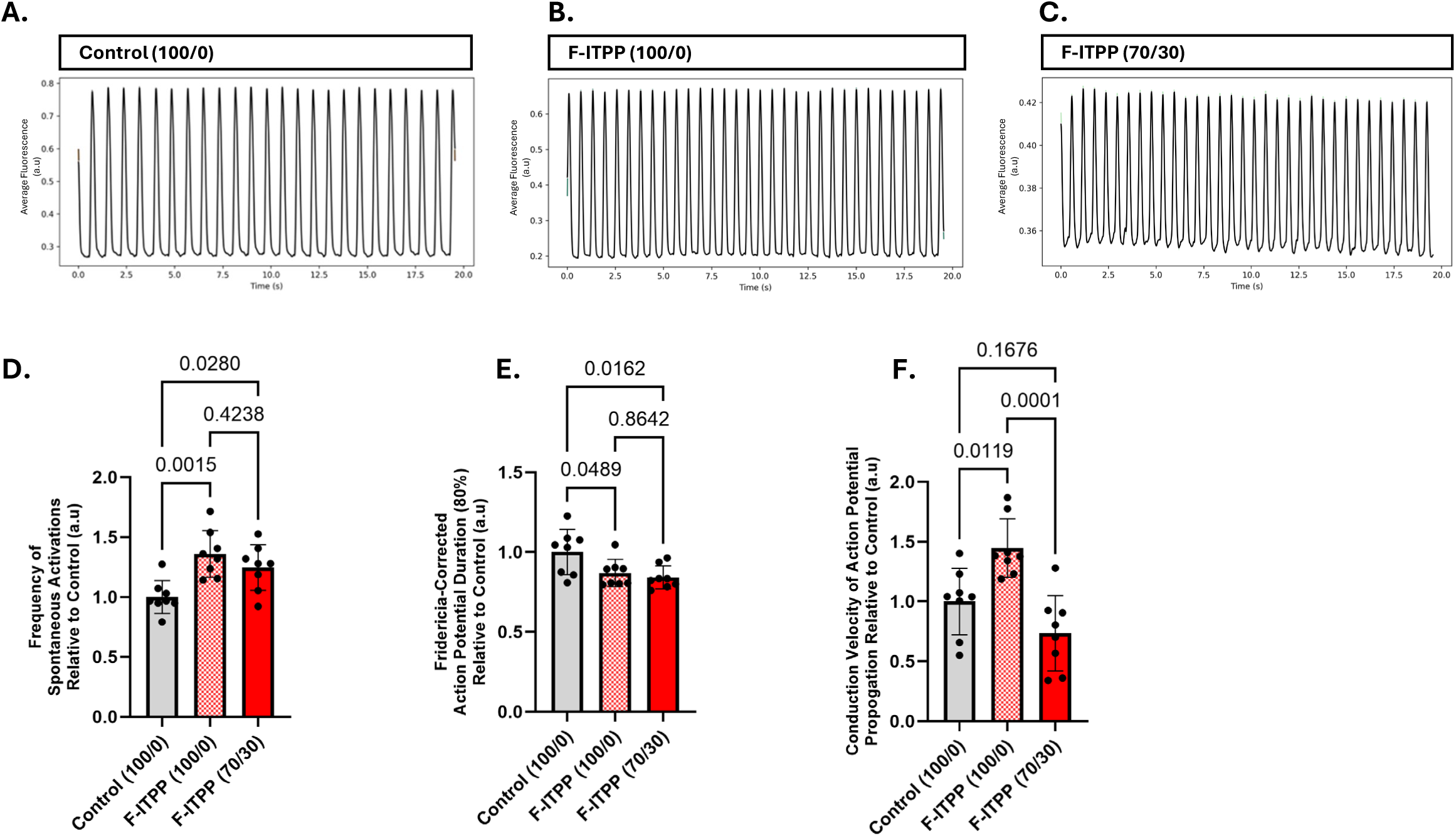
Fibrosis introduces a conduction block to intermittent tachypacing of hiPSC-aCMs. (A-C) Representative time-sequence plots of action potentials from control (A), ITPP-treated 100/0 (B), and (C) 70/30 hiPSC-aCM/haCF co-culture (F-ITPP). (D) Both 100/0 and 70/30 preparations responded to ITPP with increased frequency of spontaneous activations (F-ITPP 100/0: Δ=+36±8%, P=0.0015; F-ITPP 70/30: Δ=+25±8%, P=0.0280), and (E) shortened action potential duration (80%) (F-ITPP 100/0: Δ=-13±6%, P=0.0389; F-ITPP 70/30: Δ=-16±6%, P=0.0122). (F) F-ITPP 100/0 exhibited an increase to conduction velocity (F-ITPP 100/0: Δ=+45±13%, P=0.0089), while the F-ITPP 70/30 showed conduction slowing (F-ITPP 70/30: Δ=-27±15%; P=0.0001).

## DISCUSSION

hiPSC-CMs represent a powerful model for investigating patient-specific genetic variations and acquired diseases (24–32). Electrical pacing of hiPSC-CMs has been previously employed to improve the cardiomyocyte maturity and to induce cardiac arrhythmias. Acute and chronic effect of electric pacing was tested within a physiological range (60 beats/min) for 24 hours and 7 days respectively. Although there was no difference observed between the acute and chronic pacing regimens in relation to cell size, reactive oxygen species or apoptosis, the Ca^2+^ transient amplitude and the upstroke velocity improved in 7 days pacing with a decrease in late Na^+^ current (33). It also led to enlarged cell size with elongated phenotype (34). In a self-organizing 3-dimensional tissue ring hiPSC-CM model, rapid pacing up to 4 Hz for 2 weeks induced re-entrant waves that resulted in improved contractility, structural and Ca^2+^ handling maturation (35). In a previous study, intermittent optogenetic tachypacing of atrial hiPSC-CMs cultured into engineered heart tissue was employed with channelrhodopsin-2 transfection. The study observed that intermittent tachypacing led to an increase in action potential amplitude and a shift in the spontaneous beating pattern. However, no significant shortening of APD nor significant increase to spontaneous beat frequency was observed, phenomena commonly associated with AF remodeling (36) and another study using atrial engineered human myocardium (aEHM) and iPSC-atrial CMs subjected to 24 hours of pacing revealed AF remodeling characteristics (37). Despite demonstrating certain remodeling effects under both acute and chronic pacing regimens, these studies are limited by the use of hiPSC-CMs with an immature phenotype, which may not fully recapitulate the complex electrophysiological properties observed in mature atrial tissue during AF.

The aim of our study was to develop an improved human *in vitro* model for AF using hiPSC-aCMs by addressing challenges related to cellular maturation and chamber-specific purity observed in previous studies, as well as recapturing the fibrotic state associated with AF which will facilitate a deeper understanding of AF pathophysiology and enable more effective drug screening.

We employed tachypacing of hiPSC-aCMs outside of the human physiological range (5Hz/300bpm) to disrupt the refractory period critically. Additionally, by delivering intermittent tachypacing we alternated periods of rapid depolarizing stimuli targeting different phases of the repolarization process with periods of undisturbed action potentials. This allowed us to assess the implications of disturbances of the refractory period and normal repolarization processes. This approach facilitated the investigation of arrhythmogenic potential and syncytial behavior under stress conditions resembling atrial fibrillation. The electrophysiological findings in ITPP-treated hiPSC-aCMs were consistent with previous studies showing tachycardia-induced electrical remodeling in both humans (38) and animal models of AF (39). Specifically, ITPP treatment led to a significant increase in spontaneous AP frequency and a shortening of APD in hiPSC-aCM syncytia. However, we did not observe significant changes to the AP upstroke slope under ITPP. Interestingly, optogenetic tachypacing approaches in both intermittent and continuous protocols noted an improvement to the AP upstroke slope (36,37). Furthermore, while the faster conduction appears to serve as a compensatory mechanism in our model using pure, mature hiPSC-aCMs, it did not fully replicate the conduction slowing typically observed in clinical settings.

In addition to the observed electrophysiological changes, significant alterations in calcium handling were identified in ITPP-treated hiPSC-aCMs. These included an increased variability in spontaneous calcium release events, reduced calcium release amplitude, and slower calcium wave propagation. It is known that DADs arise from spontaneous calcium release from SR that can lead to premature action potentials hence contributing as a trigger for the onset of AF (40) which is replicated in our study. Evidence suggests that mutation in the ryanodine receptor 2 (RyR2), which is responsible for calcium release from the SR are associated with AF (41,42). Moreover, inhibition of RyR2 in Ryr2*^S2814A^* mice reduced susceptibility to AF (43). Our future studies will explore these mechanistic pathways further, alongside the roles of other critical factors in calcium handling, such as SR Ca^2+^-ATPase (SERCA) pump and the sodium-calcium exchanger (NCX) both of which play pivotal roles in calcium regulation during AF (44). Furthermore, upon application of AIT, ITPP-treated cells exhibited a reduced calcium upstroke slope and impaired reconstitution of diastolic calcium, which could contribute to conditions that predispose to calcium overload.

We co-cultured hiPSC-aCMs with aCFs for seven days to recapitulate the cues delivered to cardiomyocytes during fibrosis: namely, deposition of ECM by aCFs, activation of pro-arrhythmic signaling pathways by aCF-derived cytokines and coupling between cardiomyocytes and aCFs. A previous study has shown that patterned co-culture have a converse effect and improved structural and electrical remodeling instead of degradation of function. Additionally, this micropattern-defined co-culture was not subjected to tachypacing to mimic AF pathophysiological cues (45). We employed a co-culture system that mimicked endomysial fibrosis of AF (46). The 70/30 ratio of hiPSC-aCMs to haCFs exhibited the most pronounced fibroblast-induced changes, including a significant upregulation of collagen III and TGFβ1, both of which are known to be involved in fibrosis and the proarrhythmic substrate in AF. The electrophysiological remodeling we observed in these co-cultures were reduced AP frequency and prolonged APD, which we speculate due to the altered ion channel expression, delayed repolarization kinetics, change in resting potential due to myocyte-fibroblast coupling, where fibroblasts may contribute as a current sink (47). Further, the slowed conduction velocity is consistent with the fibrotic remodeling seen in AF, where increased extracellular matrix deposition can impair electrical conduction and lead to re-entrant circuits. Additionally, the changes in calcium handling, including prolonged CaTD_80%_, reduced calcium upstroke slope and amplitude of calcium release, suggest that fibroblast-induced alterations may contribute to the arrhythmogenic substrate in AF.

The co-culture experiments also revealed that the 90/10 ratio show a modest decrease in diastolic calcium levels was observed, suggesting that a lower fibroblast content may have subtler effects on calcium handling. In contrast, the 70/30 ratio co-cultures exhibited more pronounced alterations in calcium homeostasis, as previously mentioned. These findings underscore the importance of fibroblast-to-cardiomyocyte ratio in modulating the electrophysiological and calcium-handling properties of the cells, potentially offering insights into the influence of fibroblast proliferation in AF susceptibility.

Finally, to model advanced stages of AF, we combined the previous modules into a single model by submitting cell cultures with endomysial fibrosis characteristics to intermittent tachypacing to obtain an entourage effect of proarrhythmogenic cues experienced by AF patients. We observed that the 70/30 ratio of hiPSC-aCMs and haCFs exhibited a marked increase in the frequency of spontaneous activations and a shortening of APD. Also, a significant decrease in conduction velocity was observed, suggesting that fibroblast presence exacerbates the conduction block and electrical instability induced by tachypacing. The lack of significant changes in AP frequency and APD between the F-ITPP 70/30 and F-ITPP 100/0 co-culture conditions suggests that fibroblasts may override the ITPP effects on electrophysiological changes, but show structural remodeling coupled with conduction slowing.

Overall, the combination of ITPP and fibroblast co-culture in hiPSC-aCMs provides a robust *in vitro* model to study the electrical and calcium-handling alterations that underlie AF. The insights gained from this study may help to elucidate the AF mechanisms, offering potential targets for therapeutic intervention. This model could be further integrated into frameworks such as the Comprehensive In Vitro Proarrhythmia Assay (CiPA) and the Japan iPS Cardiac Safety Assessment (JiCSA), thereby enhancing the specificity of proarrhythmic risk assessment (48,49) and to serve as a valuable testbed for new AF treatments. Future studies are warranted to explore the precise molecular pathways driving these remodeling processes and their implications for the development of AF-targeted treatments.

### Limitations

Although this study addresses maturation of hiPSC-CMs, controlled heterogeneity, chamber specificity, and effective electrical remodelling of hiPSC-aCMs, it is important to acknowledge that other dimensions of AF such as effect of other cell types like macrophages, and sympathetic and parasympathetic neurons were not recapitulated in our *in vitro* model. As such, it partially recapitulates the intricate environmental cues and systemic regulatory mechanisms of an *in-situ* heart. Additionally, we did not intend to include in this study the inherent genetic predispositions to certain AF patients. Furthermore, while optical mapping provides a composite of actuation and concatenation of different ion channels and electronic-pumps, additional electrophysiological analysis at the single-cell level will provide refinement of understanding of electrophysiology changes present in this model. Future studies will aim to incorporate these analyses, as well as investigate the mechanistic role of AF through transcriptomics and proteomics using deep learning approaches.

### Conclusions

In conclusion, our study demonstrates that ITPP-treated mature hiPSC-aCM monolayers exhibit significant electrophysiological alterations, including increased spontaneous action potential frequency, shortened action potential duration, reduced amplitude of calcium release, slower calcium wave propagation, and impaired diastolic calcium recovery. When electrical changes are compounded with structural remodeling, reduced conduction velocity was observed, and another hallmark of AF was replicated *in vitro*.

### Clinical relevance

The observed AF phenotype, characterized by alterations in electrical activity, calcium dynamics, and structural remodeling, provides a robust *in vitro* model for investigating the underlying mechanisms of AF.

### Translational Outlook

This study presents a refined AF model that facilitates the investigation of AF pathophysiology, and serves as a platform for drug screening, potentially advancing the development of more effective AF therapeutic strategies.

## Acknowledgements

We gratefully acknowledge the technical support provided by the members at the Frankel Cardiovascular Center Regeneration Core Laboratory at the University of Michigan.

## Data availability

The data supporting the results of this study are provided within the article and its supplementary materials. Raw data are available upon reasonable request from the corresponding author.

## Abbreviations

AF: Atrial Fibrillation
ITPP: Intermittent tachypacing protocol
hiPSC-aCM: human induced-pluripotent stem cells-derived atrial cardiomyocytes
haCF: human atrial cardiac fibroblast
APD_80%_: Action potential duration at 80% repolarization
CaTD_80%_: Calcium transient duration at 80% relaxation
Col3A1: Collagen III
TGFβ1: Transforming growth factor beta 1
cTnI: cardiac troponin I
TE7: Monoclonal antibody targeting the fibroblast growth factor receptor 1 (FGFR1) antigen
AIT: Arrythmia inducibility test

## REFERENCES

1. Schuttler D, Bapat A, Kaab S et al. Animal Models of Atrial Fibrillation. Circ Res 2020;127:91–110.

2. Yamamoto C, Trayanova NA. Atrial fibrillation: Insights from animal models, computational modeling, and clinical studies. EBioMedicine 2022;85:104310.

3. Heijman J, Sutanto H, Crijns H, Nattel S, Trayanova NA. Computational models of atrial fibrillation: achievements, challenges, and perspectives for improving clinical care. Cardiovasc Res 2021;117:1682–1699.

4. Nattel S, Sager PT, Huser J, Heijman J, Dobrev D. Why translation from basic discoveries to clinical applications is so difficult for atrial fibrillation and possible approaches to improving it. Cardiovasc Res 2021;117:1616–1631.

5. Sayed N, Ameen M, Wu JC. Personalized medicine in cardio-oncology: the role of induced pluripotent stem cell. Cardiovasc Res 2019;115:949–959.

6. Zhang L, Tian L, Dai X et al. Pluripotent stem cell-derived CAR-macrophage cells with antigen-dependent anti-cancer cell functions. J Hematol Oncol 2020;13:153.

7. Kuhl L, Graichen P, von Daacke N et al. Human Lung Organoids-A Novel Experimental and Precision Medicine Approach. Cells 2023;12.

8. Vatine GD, Barrile R, Workman MJ et al. Human iPSC-Derived Blood-Brain Barrier Chips Enable Disease Modeling and Personalized Medicine Applications. Cell Stem Cell 2019;24:995–1005 e6.

9. Wang S, Du Y, Zhang B et al. Transplantation of chemically induced pluripotent stem-cell-derived islets under abdominal anterior rectus sheath in a type 1 diabetes patient. Cell 2024;187:6152–6164 e18.

10. Arjmand B, Goodarzi P, Mohamadi-Jahani F, Falahzadeh K, Larijani B. Personalized Regenerative Medicine. Acta Med Iran 2017;55:144–149.

11. Parrotta EI, Lucchino V, Scaramuzzino L, Scalise S, Cuda G. Modeling Cardiac Disease Mechanisms Using Induced Pluripotent Stem Cell-Derived Cardiomyocytes: Progress, Promises and Challenges. Int J Mol Sci 2020;21.

12. Desai D, Song T, Singh RR et al. MYBPC3 D389V Variant Induces Hypercontractility in Cardiac Organoids. Cells 2024;13.

13. Maurissen TL, Kawatou M, Lopez-Davila V, Minatoya K, Yamashita JK, Woltjen K. Modeling mutation-specific arrhythmogenic phenotypes in isogenic human iPSC-derived cardiac tissues. Sci Rep 2024;14:2586.

14. Duboscq-Bidot L, Hoareau B, Ader F, Fontaine V, Villard E. Generation of CRISPR-Cas9 edited human induced pluripotent stem cell line carrying BAG3 V468M mutation in its BAG domain. Stem Cell Res 2024;74:103294.

15. Benzoni P, Campostrini G, Landi S et al. Human iPSC modelling of a familial form of atrial fibrillation reveals a gain of function of If and ICaL in patient-derived cardiomyocytes. Cardiovasc Res 2020;116:1147–1160.

16. Block T, Creech J, da Rocha AM et al. Human perinatal stem cell derived extracellular matrix enables rapid maturation of hiPSC-CM structural and functional phenotypes. Sci Rep 2020;10:19071.

17. Allan A, Creech J, Hausner C et al. High-throughput longitudinal electrophysiology screening of mature chamber-specific hiPSC-CMs using optical mapping. iScience 2023;26:107142.

18. Burridge PW, Matsa E, Shukla P et al. Chemically defined generation of human cardiomyocytes. Nat Methods 2014;11:855–60.

19. Biendarra-Tiegs SM, Li X, Ye D, Brandt EB, Ackerman MJ, Nelson TJ. Single-Cell RNA-Sequencing and Optical Electrophysiology of Human Induced Pluripotent Stem Cell-Derived Cardiomyocytes Reveal Discordance Between Cardiac Subtype-Associated Gene Expression Patterns and Electrophysiological Phenotypes. Stem Cells Dev 2019;28:659–673.

20. Kleinsorge M, Cyganek L. Subtype-Directed Differentiation of Human iPSCs into Atrial and Ventricular Cardiomyocytes. STAR Protoc 2020;1:100026.

21. Cyganek L, Tiburcy M, Sekeres K et al. Deep phenotyping of human induced pluripotent stem cell-derived atrial and ventricular cardiomyocytes. JCI Insight 2018;3.

22. Monteiro da Rocha A, Allan A, Block T, Creech J, Herron TJ. High-Throughput Cardiotoxicity Screening Using Mature Human Induced Pluripotent Stem Cell-Derived Cardiomyocyte Monolayers. J Vis Exp 2023.

23. Zhang J, Tao R, Campbell KF et al. Functional cardiac fibroblasts derived from human pluripotent stem cells via second heart field progenitors. Nat Commun 2019;10:2238.

24. Ye L, Chang JC, Lin C, Sun X, Yu J, Kan YW. Induced pluripotent stem cells offer new approach to therapy in thalassemia and sickle cell anemia and option in prenatal diagnosis in genetic diseases. Proc Natl Acad Sci U S A 2009;106:9826–30.

25. Urbach A, Bar-Nur O, Daley GQ, Benvenisty N. Differential modeling of fragile X syndrome by human embryonic stem cells and induced pluripotent stem cells. Cell Stem Cell 2010;6:407–11.

26. Itzhaki I, Maizels L, Huber I et al. Modelling the long QT syndrome with induced pluripotent stem cells. Nature 2011;471:225–9.

27. Narsinh K, Narsinh KH, Wu JC. Derivation of human induced pluripotent stem cells for cardiovascular disease modeling. Circ Res 2011;108:1146–56.

28. Kudva YC, Ohmine S, Greder LV et al. Transgene-free disease-specific induced pluripotent stem cells from patients with type 1 and type 2 diabetes. Stem Cells Transl Med 2012;1:451–61.

29. Teo AK, Windmueller R, Johansson BB et al. Derivation of human induced pluripotent stem cells from patients with maturity onset diabetes of the young. J Biol Chem 2013;288:5353–6.

30. Shang L, Hua H, Foo K et al. beta-cell dysfunction due to increased ER stress in a stem cell model of Wolfram syndrome. Diabetes 2014;63:923–33.

31. Simsek S, Zhou T, Robinson CL et al. Modeling Cystic Fibrosis Using Pluripotent Stem Cell-Derived Human Pancreatic Ductal Epithelial Cells. Stem Cells Transl Med 2016;5:572–9.

32. Laperle AH, Sances S, Yucer N et al. iPSC modeling of young-onset Parkinson’s disease reveals a molecular signature of disease and novel therapeutic candidates. Nat Med 2020;26:289–299.

33. Knierim M, Bommer T, Paulus M et al. Cellular calcium handling and electrophysiology are modulated by chronic physiological pacing in human induced pluripotent stem cell-derived cardiomyocytes. Am J Physiol Heart Circ Physiol 2024;327:H1244–H1254.

34. Zhou J, Cui B, Wang X et al. Overexpression of KCNJ2 enhances maturation of human-induced pluripotent stem cell-derived cardiomyocytes. Stem Cell Res Ther 2023;14:92.

35. Li J, Zhang L, Yu L et al. Circulating re-entrant waves promote maturation of hiPSC-derived cardiomyocytes in self-organized tissue ring. Commun Biol 2020;3:122.

36. Lemoine MD, Lemme M, Ulmer BM et al. Intermittent Optogenetic Tachypacing of Atrial Engineered Heart Tissue Induces Only Limited Electrical Remodelling. J Cardiovasc Pharmacol 2020;77:291–299.

37. Seibertz F, Rubio T, Springer R et al. Atrial fibrillation-associated electrical remodelling in human induced pluripotent stem cell-derived atrial cardiomyocytes: a novel pathway for antiarrhythmic therapy development. Cardiovasc Res 2023;119:2623–2637.

38. Nattel S, Burstein B, Dobrev D. Atrial remodeling and atrial fibrillation: mechanisms and implications. Circ Arrhythm Electrophysiol 2008;1:62–73.

39. Nattel S, Shiroshita-Takeshita A, Brundel BJ, Rivard L. Mechanisms of atrial fibrillation: lessons from animal models. Prog Cardiovasc Dis 2005;48:9–28.

40. Denham NC, Pearman CM, Caldwell JL et al. Calcium in the Pathophysiology of Atrial Fibrillation and Heart Failure. Front Physiol 2018;9:1380.

41. Shan J, Xie W, Betzenhauser M et al. Calcium leak through ryanodine receptors leads to atrial fibrillation in 3 mouse models of catecholaminergic polymorphic ventricular tachycardia. Circ Res 2012;111:708–17.

42. Kushnir A, Wajsberg B, Marks AR. Ryanodine receptor dysfunction in human disorders. Biochim Biophys Acta Mol Cell Res 2018;1865:1687–1697.

43. Chelu MG, Sarma S, Sood S et al. Calmodulin kinase II-mediated sarcoplasmic reticulum Ca2+ leak promotes atrial fibrillation in mice. J Clin Invest 2009;119:1940–51.

44. Yeh YH, Wakili R, Qi XY et al. Calcium-handling abnormalities underlying atrial arrhythmogenesis and contractile dysfunction in dogs with congestive heart failure. Circ Arrhythm Electrophysiol 2008;1:93–102.

45. Brown GE, Han YD, Michell AR et al. Engineered cocultures of iPSC-derived atrial cardiomyocytes and atrial fibroblasts for modeling atrial fibrillation. Sci Adv 2024;10:eadg1222.

46. Maesen B, Verheule S, Zeemering S et al. Endomysial fibrosis, rather than overall connective tissue content, is the main determinant of conduction disturbances in human atrial fibrillation. Europace 2022;24:1015–1024.

47. Jacquemet V, Henriquez CS. Modelling cardiac fibroblasts: interactions with myocytes and their impact on impulse propagation. Europace 2007;9 Suppl 6:vi29–37.

48. Blinova K, Dang Q, Millard D et al. International Multisite Study of Human-Induced Pluripotent Stem Cell-Derived Cardiomyocytes for Drug Proarrhythmic Potential Assessment. Cell Rep 2018;24:3582–3592.

49. Kanda Y, Yamazaki D, Osada T, Yoshinaga T, Sawada K. Development of torsadogenic risk assessment using human induced pluripotent stem cell-derived cardiomyocytes: Japan iPS Cardiac Safety Assessment (JiCSA) update. J Pharmacol Sci 2018;138:233–239.

