## Supplementary Material for "Modeling Atrial Fibrillation through Intermittent Tachypacing-Induced Remodeling in hiPSC-Derived Atrial Cardiomyocytes and Atrial Fibroblast"

### METHODS

#### *Atrial Cardiac Myocyte Differentiation of hiPSCs*

Human iPSCs were cultured on multi-well plates coated with Matrigel (Corning) and maintained in either StemMACS iPS-Brew XF (Miltenyi Biotec) or mTESR Plus (STEMCELL Technologies) media. hiPSCs were differentiated an into an atrial-specific lineage using a modified GiWi protocol (1). After the cells reach 90-100% confluency, on differentiation (day 0), Basal Medium (RPMI 1640 supplemented with L-Glutamine and 25 mM HEPES (Gibco), supplemented with 0.5 mg/mL BSA (Sigma), 0.2 mg/mL L-Ascorbic Acid (Sigma), and 4 μM CHIR99021 (Selleck) was added. 48 hours following CHIR99021 treatment (day 2), medium was changed to Basal Medium containing 5 μM IWP4 (Selleck). On day 3, medium was switched to Basal Medium containing 5 μM IWP4 and 1 μM Retinoic Acid (Selleck). On day 4, the medium was changed with Basal Medium containing 1 μM Retinoic Acid. On day 6, the medium was changed to Basal Medium without any small molecules. On day 8, the medium was switched to RPMI 1640 with L-Glutamine (Gibco) supplemented with B-27 (Gibco). After day 8, the medium was refreshed every 48-72 hours with RPMI 1640 with L-Glutamine supplemented with B-27 until cell purification.

#### *Purification and maturation of hiPSC-aCMs*

Human iPSC-derived atrial differentiations were purified by negative-selection using biotin-conjugated magnetic bead sorting with an autoMACS Pro Separator (Miltenyi). The cells were dissociated with 0.25% Trypsin/EDTA (Fisher). Following trypsinization, EB20 Medium (80% DMEM/F12, 0.1 mmol/L nonessential amino acids, 1 mmol/L L-glutamine, 0.1 mmol/L β-mercaptoethanol, 20% FBS and 10 μmol/L blebbistatin) with 1 μM ROCK inhibitor (ROCKi; Y-27632, Tocris) was added. The cells were dissociated and then strained through a 70 μm cell strainer. Total cell count was estimated using a hemocytometer. The cell suspension was centrifuged at 1000 RPM for 5 minutes and the supernatant was aspirated. The cell pellet was resuspended in cold autoMACS Running Buffer (Miltenyi) and centrifuged again at 1000 RPM for 5 minutes. The cells were purified by depleting iPSC-derived non-cardiomyocytes by magnetic separation using Miltenyi isolation kit according to the manufacturer's instruction. Maturation of hiPSC-aCMs on MatrixPlus was facilitated as previously described (2-4). A 96-well black microclear plate was coated with MatrixPlus, then rehydrated with HBSS with calcium and magnesium at 37 °C for 30 mins. The purified hiPSC-derived atrial cardiomyocytes were replated at a density of 95,000 cells/well. The hiPSC-aCMs were initially cultured in EB20 Medium with ROCKi for 48 hours, after which the medium was replaced with RPMI 1640 with L-Glutamine, supplemented with B-27 for 5 days, with media changes every 48 hours. All experiments were initiated 7 days post-purification, at which point hiPSC-aCMs had formed mature syncytia.

***Culture of haCFs***

Cryopreserved human donor atrial cardiac fibroblasts (haCFs; NHCF-A Human Cardiac Atrial Fibroblasts; Lonza, #CC-2903) were obtained from Lonza Biosciences (Walkersville, MD) and thawed into FBM$^{TM}$ Basal Medium (Lonza, #CC-3131) supplemented with FGM$^{TM}$-3 SingleQuot Supplements (Lonza, #CC-4525). The cells were cultured in T-25 cell culture flasks coated with 0.1% gelatin (Type A, Sigma) in 1X PBS. The haCFs were maintained for 2-4 days until they reached 70-80% confluency, at which point they were passaged for experiments (not more than 5 passages) according to the manufacturer's protocol.

***Genetically Encoded Calcium Indicator Transfection of hiPSC-aCMs***

To enable long-term optical mapping of calcium transients, hiPSC-aCMs submitted to ITPP were transfected with the genetically encoded calcium indicator, GCaMP6 fast variant, as previously described (3). Recombinant adenoviruses (AdGCaMP6f, Vector Biolabs, Malvern, PA, #1910) were prepared and had a titer of $1 \times 10^{10}$ PFU/mL. Human iPSC-aCMs were transfected with a load of 5 multiplicity of infection (MOI) GCaMP6f in 100 μL of RPMI with L-Glutamine and without phenol red supplemented with B-27 (Gibco) for 48 hours. Cells were further incubated in RPMI with L-Glutamine and without phenol red for 3 days to allow for endogenous expression of GCaMP6f; all transfection protocols were performed within a 7-day window prior to induction of ITPP.

***Arrhythmia Inducibility Test***

Following 7 days of intermittent-tachypacing, hiPSC-aCM monolayers were subjected to a point stimulation test with simultaneous optical mapping. Optical mapping of calcium transients

(CaTs) was conducted by placing the 8-well plate beneath a high-speed CCD camera (200 fps, 80 x 80 pixels; Red-Shirt Little Joe, Scimeasure, Decatur, GA). A heating block was used to maintain physiological temperature (37 °C) during recordings. An emission filter (515 nm; Chroma) appropriate to the loaded GCaMP6f, was used in combination with blue LED illumination to capture spontaneous or electrically induced CaTs. Point stimulation of the monolayers was facilitated using a dual-pronged electrode coupled to a MyoPacer$^{TM}$ power bank (IONOptix). The electrode was placed in direct contact with the monolayers under the guidance from the CCD camera and the monolayers were paced at 3.5 Hz (25 V) for 30s. Optical mapping of CaTs began 5s prior to termination of point stimulation, and resultant spontaneous CaTs were then recorded for 15s. Data analysis was conducted with a custom software (Scroll5).

#### *Immunocytochemistry*

Cells were fixed with 4% paraformaldehyde for 20 minutes and permeabilized in 0.1% Triton-X100 (Fisher) for 10 minutes at room temperature. After permeabilization, cells were washed once with 1x PBS and incubated for 30 minutes at room temperature in a blocking solution of 1% normal donkey serum in 1x PBS containing 0.1% Triton-X100. Primary antibodies (Table 1) were diluted in the blocking solution and incubated with the cells for 1 hour at room temperature. Cells were then washed with 1x PBS twice. Secondary antibodies (Table 1) were then applied in the blocking solution and incubated for 30 minutes at room temperature in the dark. Cells were washed with 1x PBS prior to nucleic acid staining with DAPI in 1x PBS (1:1000) for 5 minutes. Additional washes with 1x PBS were performed before imaging. Imaging was performed using either a Cytation 5 Imaging Reader (Biotek), Incucyte SX5 (Sartorius), or a Nikon A1R confocal microscope (Nikon).

#### *Statistical Analysis*

All data are presented as mean ± standard deviation (SD) and normalized to the corresponding control values. Mean differences (Δ) between groups are reported along with the standard error of the difference. Outlier analysis was performed using ROUT with 1% sensitivity. Outliers between interrelated measurements (dF/F0, F0, Upstroke Slope, Conduction Velocity; or Frequency, CaTD80 / APD80, CaTDTri / APDTri) were removed. For comparison of 2 groups, Kolmogorov-Smirnov test of normality was performed with subsequent unpaired parametric T-Test or Mann-Whitney U Test. For comparison of >2 groups, 1-Way-ANOVA was performed with Bonferroni comparison of individual groups to control, or Tukey's test for multiple comparisons between groups. A 2-tailed $P<0.05$ indicated statistical significance.

**Table 1: List of antibodies utilized in this study**

| Antibody | Company | Catalog # | Species | Dilution |
|---|---|---|---|---|
| **Primary antibodies:** | | | | |
| **Anti-TE-7 (ER-TR7)** | Millipore | CBL271 | Mouse | 1:100 |
| **Anti-α-Actinin** | Sigma | A7811 | Mouse | 1:500 |
| **Anti-cTnT** | Abcam | 45932 | Rabbit | 1μg/mL |
| **Anti-TGFβ1** | Santa Cruz | sc-146 | Rabbit | 1:100 |
| **Anti-Collagen III** | Invitrogen | PA5-34787 | Rabbit | 1:500 |
| **Secondary antibodies:** | | | | |
| **Donkey anti-Rabbit (594)** | Thermo Fisher | A21207 | Anti-Rabbit | 1:500 |
| **Donkey anti-mouse (488)** | Jackson ImmunoResearch | 715-545-150 | Anti-Mouse | 1:500 |

### REFERENCES


1. Cyganek L, Tiburcy M, Sekeres K et al. Deep phenotyping of human induced pluripotent stem cell-derived atrial and ventricular cardiomyocytes. JCI Insight 2018;3.

2. Block T, Creech J, da Rocha AM et al. Human perinatal stem cell derived extracellular matrix enables rapid maturation of hiPSC-CM structural and functional phenotypes. Sci Rep 2020;10:19071.

3. Allan A, Creech J, Hausner C et al. High-throughput longitudinal electrophysiology screening of mature chamber-specific hiPSC-CMs using optical mapping. iScience 2023;26:107142.

4. Monteiro da Rocha A, Allan A, Block T, Creech J, Herron TJ. High-Throughput Cardiotoxicity Screening Using Mature Human Induced Pluripotent Stem Cell-Derived Cardiomyocyte Monolayers. J Vis Exp 2023.
